# Bisulfite-free, Base-resolution, and Quantitative Sequencing of Cytosine Modifications

**DOI:** 10.1101/307538

**Authors:** Yibin Liu, Paulina Siejka, Gergana Velikova, Fang Yuan, Marketa Tomkova, Chunsen Bai, Lei Chen, Ying Bi, Benjamin Schuster-Boeckler, Chun-Xiao Song

**Author notes:** P.S. and G.V. contributed equally to this work. Corresponding authors. (B.S.-B); (C.-X.S.).

## Abstract

The deamination of unmodified cytosine to uracil by treatment with bisulfite has for decades been the gold standard for sequencing epigenetic DNA modifications including 5-methylcytosine (5mC) and 5-hydroxymethylcytosine (5hmC). However, this harsh chemical reaction degrades the majority of the DNA and generates sequencing libraries with low complexity. Here, we present a novel bisulfite-free and base-resolution sequencing method, TET Assisted Pic-borane Sequencing (TAPS), for detection of 5mC and 5hmC. TAPS relies on mild reactions, detects modifications directly without affecting unmodified cytosines and can be adopted to detect other cytosine modifications. Compared with bisulfite sequencing, TAPS results in higher mapping rates, more even coverage and lower sequencing costs, enabling higher quality, more comprehensive and cheaper methylome analyses.

**One Sentence Summary:** A bisulfite-free and base-resolution method to directly sequence epigenetically modified cytosine.

---

DNA cytosine modifications are important epigenetic mechanisms that play crucial roles in a broad range of biological processes from gene regulation to normal development *(1)*. 5-Methylcytosine (5mC) and 5-hydroxymethylcytosine (5hmC) are by far the two most common epigenetic marks found in the mammalian genome. 5hmC is generated from 5mC by the ten-eleven translocation (TET) family of dioxygenases *(2)*. TET can further oxidize 5hmC to 5-formylcytosine (5fC) and 5-carboxylcytosine (5caC), which exist in much lower abundances in the mammalian genome compared to 5mC and 5hmC (10-fold to 100-fold lower than that of 5hmC) *(3)*. Aberrant DNA methylation and hydroxymethylation have been associated with various diseases and are well-accepted hallmarks of cancer *(4, 5)*. Therefore, the determination of the genomic distribution of 5mC and 5hmC is not only important for our understanding of development and homeostasis, but is also invaluable for clinical applications *(6, 7)*.

The current gold standard and the only option for base-resolution and quantitative DNA methylation and hydroxymethylation analysis is bisulfite sequencing (BS) *(8, 9)*, and its derived methods including TET-assisted bisulfite sequencing (TAB-Seq) *(10)* and oxidative bisulfite sequencing (oxBS) *(11)*. All these methods employ bisulfite treatment to convert unmethylated cytosine to uracil while leaving 5mC and/or 5hmC intact. Since PCR amplification of the bisulfite-treated DNA reads uracil as thymine, the modification of each cytosine can be inferred at single base resolution, where C-to-T transitions provide the locations of the unmethylated cytosines. There are, however, two main drawbacks to bisulfite sequencing. Firstly, bisulfite treatment is a harsh chemical reaction which degrades up to 99% of the DNA due to depyrimidination under the required acidic and thermal conditions *(12)*. This severely limits its utility if sample DNA quantities are low. Secondly, bisulfite sequencing relies on the complete conversion of unmodified cytosine to thymine. Unmodified cytosine accounts for approx. 95% of the total cytosine in the human genome. Converting all these positions to thymine severely reduces sequence complexity, leading to poor sequencing quality, low mapping rates, uneven genome coverage and increased sequencing cost. Consequently, bisulfite sequencing suffers from pronounced sequencing biases and overestimation of methylation levels due to selective and context-specific DNA degradation *(13)*. To solve these problems, a mild reaction that can directly detect modified cytosine (5mC and 5hmC) at base-resolution, without affecting unmodified cytosine, is desired to accurately estimate methylation levels.

Recently, an elegant bisulfite-free and base-resolution method for sequencing 5fC has been developed based on Friedländer synthesis reaction, which can induce a 5fC-to-T transition *(14, 15)*. However, this method has limited application since 5fC is a rare modification and there is no way to convert 5mC efficiently and completely to 5fC *(16)*. There is, however, a convenient way to convert 5mC and 5hmC to 5caC. The TET enzymes readily oxidize 5mC and 5hmC to the final oxidation product 5caC *in vitro (3, 17)*. We envisioned that if we could induce a 5caC-to-T transition, it could be combined with TET oxidation of 5mC and 5hmC to enable direct detection of 5mC and 5hmC. Here we present such a 5caC-to-T transition chemistry, and its application for whole-genome base-resolution detection of cytosine modifications.

We started with a 11mer 5caC-containing DNA oligo as a model DNA, which we used to screen chemicals that could react with 5caC, as monitored by matrix-assisted laser desorption/ionization mass spectroscopy (MALDI). We discovered that certain borane-containing compounds could efficiently react with the 5caC oligo, resulting in a molecular weight reduction of 41 Da (fig. S1A and Fig. 1A). We chose 2-picoline borane (pic-borane) to further study as it is a commercially available and environmentally benign reducing agent *(18)*.

**Fig. 1.**
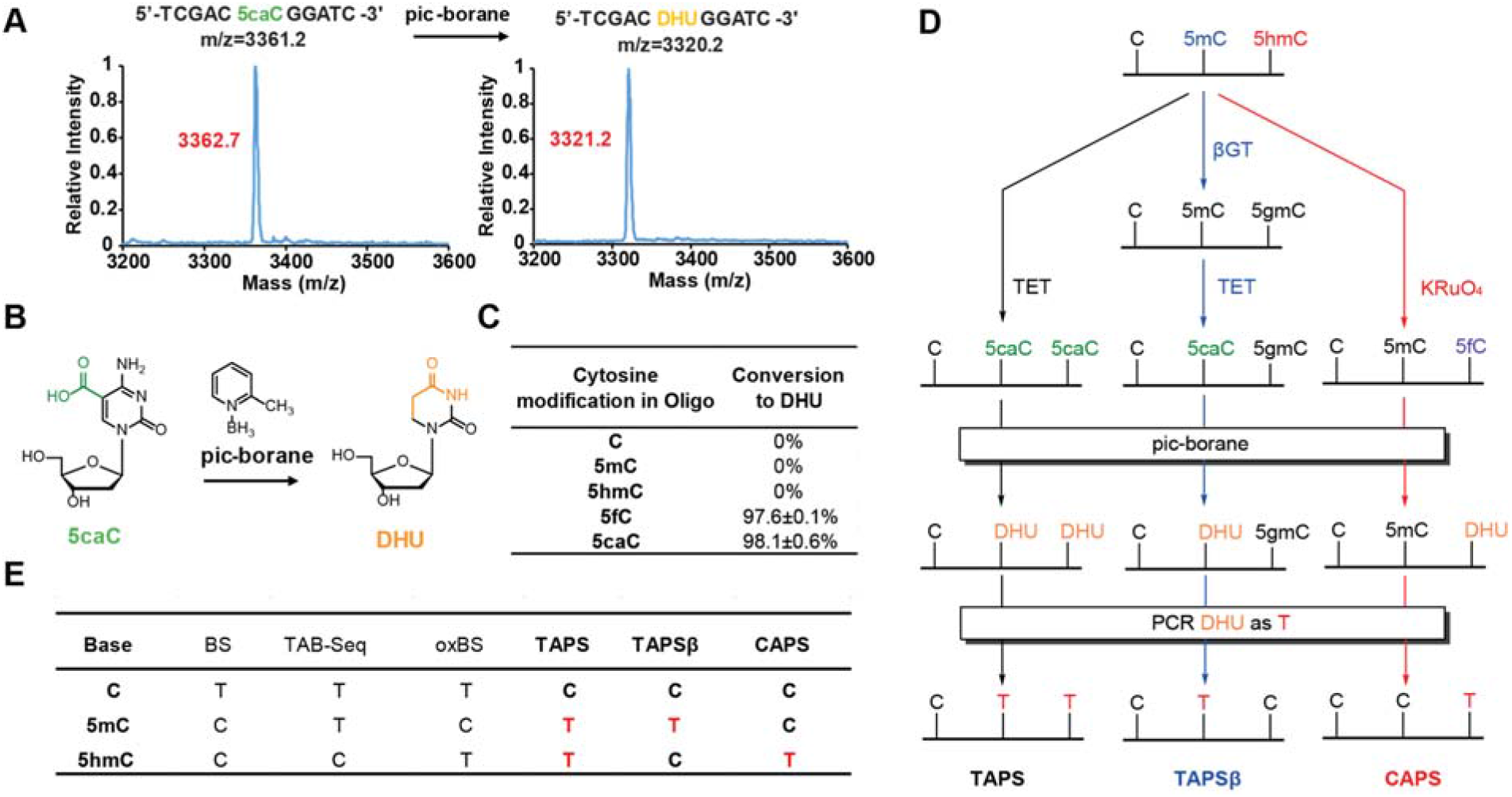
Pic-borane reaction on DNA oligos. **(A)** MALDI characterization of 5caC-containing 11mer model DNA treated with pic-borane. Calculated mass shown in black, observed mass shown in red. **(B)** Reaction of pic-borane conversion of 5caC to DHU. **(C)** The conversion rates of dC and various cytosine derivatives quantified by HPLC-MS/MS. Data shown as mean ± SD of three replicates. **(D)** Overview of the TAPS, TAPSβ, and CAPS. **(E)** Comparison of BS and related methods versus TAPS and CAPS for 5mC and 5hmC sequencing.

We repeated the reaction on 5caC single nucleoside and confirmed that pic-borane converts 5caC to dihydrouracil (DHU) (Fig. 1B and Supplementary Text). To the best of our knowledge, this is a previously unknown reductive decarboxylation/deamination reaction (fig. S1B). Interestingly, we found pic-borane can also convert 5fC to DHU through an apparent reductive deformylation/deamination mechanism (fig. S2 and S3). The detailed mechanism of both reactions remains to be defined. Quantitative analysis of the pic-borane reaction on the DNA oligo by HPLC-MS/MS confirms that pic-borane converts 5caC and 5fC to DHU with around 98% efficiency and has no activity against unmethylated cytosine, 5mC or 5hmC (Fig. 1C).

As a uracil derivative, DHU can be recognized by both DNA and RNA polymerases as thymine *(19)*. Therefore, pic-borane could induce both 5caC-to-T and 5fC-to-T transitions, and can be used for base-resolution sequencing of 5fC and 5caC, which we termed Pic-borane Sequencing (PS) (Table S1). The reaction of 5fC and 5caC with pic-borane can be blocked by hydroxylamine conjugation *(20)* and EDC coupling *(21)*, respectively (fig. S3), which allows PS to be used to sequence 5fC or 5caC specifically (Table S1). More importantly, we can now use TET enzymes to oxidize 5mC and 5hmC to 5caC, and then subject 5caC to pic-borane treatment in a process we call TET-Assisted Pic-borane Sequencing (TAPS) (Fig. 1D-E). TAPS can induce a C-to-T transition of 5mC and 5hmC, and therefore can be used for base-resolution detection of 5mC and 5hmC. Furthermore, ß-glucosyltransferase (ßGT) can label 5hmC with glucose and thereby protect it from TET oxidation *(10)* and pic-borane reduction (fig. S4), enabling the selective sequencing of only 5mC, in a process we call TAPSβ (Fig. 1D-E). 5hmC sites can then be deduced by subtraction of TAPSβ from TAPS measurements. Alternatively, we can use potassium perruthenate (KRuO4), a reagent previously used in oxidative bisulfite sequencing (oxBS) *(11)*, to replace TET as a chemical oxidant to specifically oxidize 5hmC to 5fC (fig. S4).

This approach, which we call Chemical-Assisted Pic-borane Sequencing (CAPS), can be used to sequence 5hmC specifically (Fig. 1D-E). Therefore, the PS and related methods can in principle offer a comprehensive suite to sequence all four cytosine epigenetic modifications (Fig. 1D-E, Table S1).

We next aimed to evaluate the performance of TAPS in comparison with bisulfite sequencing, the current standard and most widely used method for base-resolution mapping of 5mC and 5hmC. We used *Naegleria* TET-like oxygenase (NgTET1) since it can efficiently oxidize 5mC to 5caC *in vitro* and can be easily produced recombinantly from *E. coli (22)*. Other TET proteins such as mouse Tet1 (mTet1) can also be used *(10)*. To confirm the 5mC-to-T transition, we applied TAPS to a 222 bp model DNA containing five fully methylated CpG sites and showed that it can effectively convert 5mC to T, as demonstrated by restriction enzyme digestion (fig. S5) and Sanger sequencing (Fig. 2A). Both TET oxidation and pic-borane reduction are mild reactions, with no notable DNA degradation compared to bisulfite (fig. S6). DHU is close to a natural base, it is compatible with common DNA polymerases such as Taq DNA Polymerase and KAPA HiFi Uracil+ DNA Polymerase (fig. S7 and S8). We next applied TAPS to genomic DNA (gDNA) from mouse embryonic stem cells (mESCs). HPLC-MS/MS quantification showed that, as expected, 5mC accounts for 98.5% of cytosine modifications in the mESCs gDNA; the remainder is composed of 5hmC (1.5%) and trace amounts 5fC and 5caC, and no DHU (Fig. 2B). After NgTET1 oxidation, about 96% of cytosine modifications were oxidized to 5caC and 3% were oxidized to 5fC (Fig. 2B). After pic-borane reduction, over 99% of the cytosine modifications were converted into DHU (Fig. 2B). These results demonstrate both NgTET1 oxidation and pic-borane reduction work efficiently on genomic DNA.

**Fig. 2.**
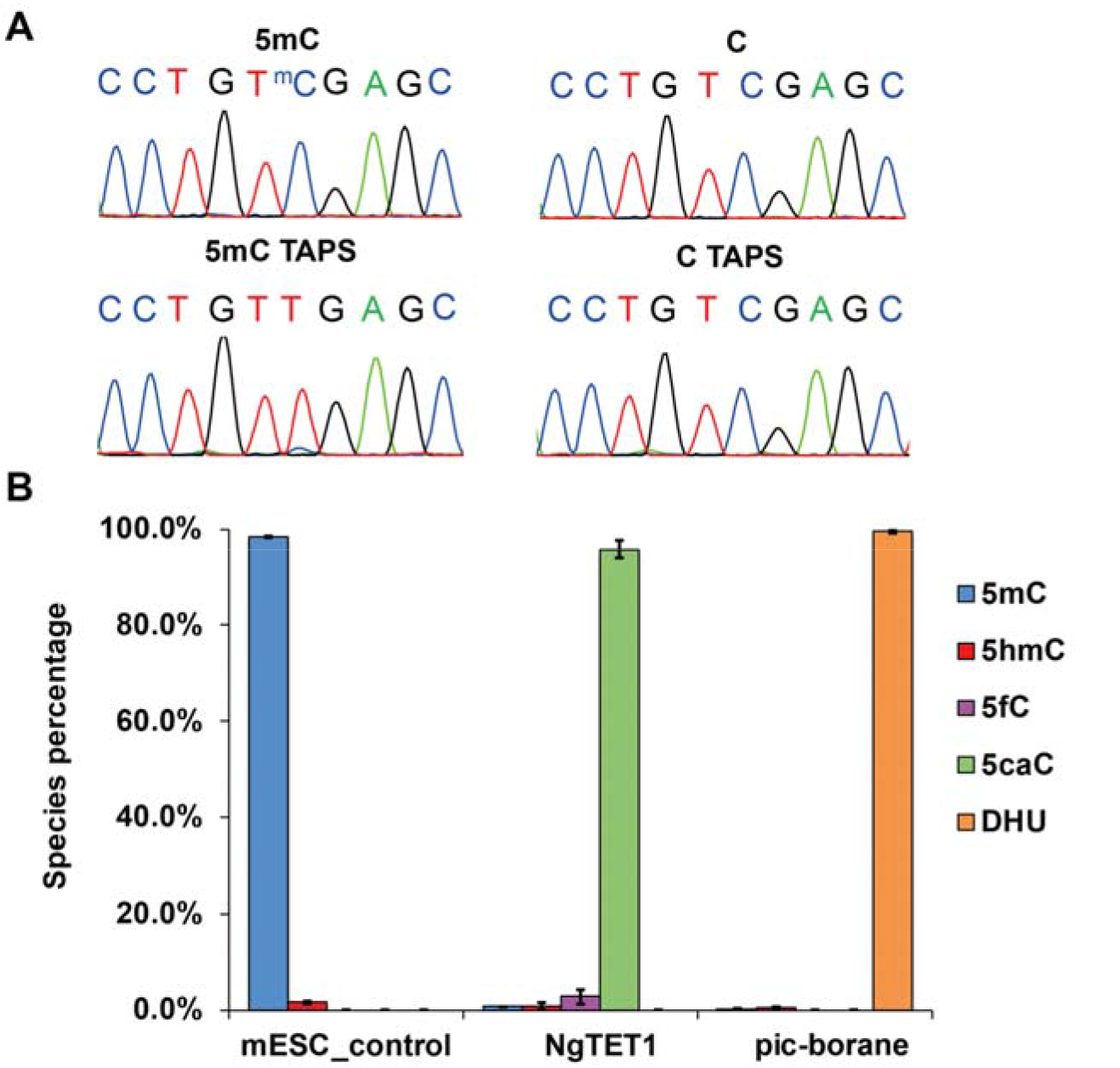
TAPS on a 222 bp model DNA and mESC gDNA. **(A)** Sanger sequencing results for the 222 bp model DNA containing 5 fully methylated CpG sites and its unmethylated control before (top) and after (bottom) TAPS. Only 5mC is converted to T after TAPS. **(B)** HPLC-MS/MS quantification of relative modification levels in the mESCs gDNA control, after NgTET1 oxidation and after pic-borane reduction. Data shown as mean ± SD of three replicates.

We subsequently performed whole genome sequencing of two samples of mESC gDNA, one converted using TAPS and the other using standard whole-genome bisulfite sequencing (WGBS) for comparison. To assess the accuracy of TAPS, we added spike-ins of different lengths that were either fully unmodified, *in vitro* methylated using CpG Methyltransferase (M.SssI) or GpC Methyltransferase (M.CviPI) (see Materials and Methods). For short spike-ins (120mer-1 and 120mer-2) containing 5mC and 5hmC, near complete conversion was observed for both modifications on both strands in both CpG and non-CpG contexts (fig. S9). Based on longer spike-ins (lambda DNA and 2kb amplicons, see Materials and Methods), the 5mC conversion rate was estimated at 85.7% for CpG and 71.5% for GpC (Fig. 3A), suggesting slightly lower conversion of TAPS in non-CpG methylation, which is consistent with the lower activity of TET proteins in non-CpG contexts *(22)*. The false positive rate (converted cytosine in unmodified spike-ins) was estimated to be below 2% (1.8% and 1.6% for CpG and GpC, respectively; Fig. 3A).

**Fig. 3.**
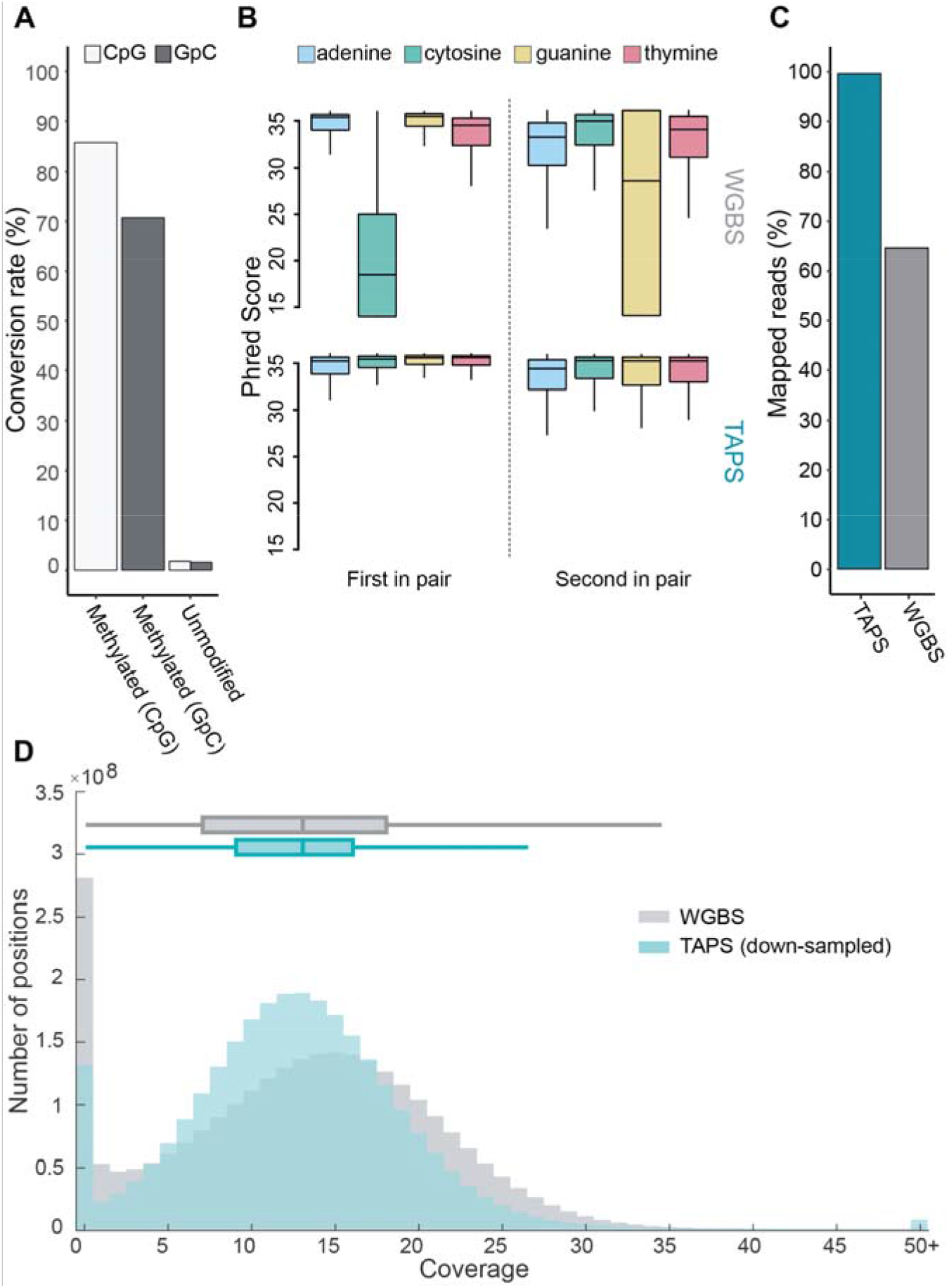
Improved sequencing quality of TAPS over WGBS. **(A)** Fraction of C in CpG or GpC converted to T on three spike-ins with different modifications. Left: lambda DNA fully methylated *in vitro* at all CpG sites. Middle: 2kb amplicon methylated at all GpC sites. Right: unmethylated 2kb amplicon. **(B)** Sequencing quality scores per base for the first and second reads in all sequenced read pairs, as reported by Illumina BaseSpace. Top: WGBS. Bottom: TAPS. Nucleotide is denoted by color. **(C)** Fraction of all sequenced read pairs (after trimming) mapped to the genome. **(D)** Comparison of coverage across all bases of the mouse genome between WGBS and TAPS. To account for differences in sequencing depth, all mapped TAPS reads were down-sampled to match the mean coverage of WGBS across the genome. Positions with coverage above 50× are shown in the last bin.

Due to the conversion of nearly all cytosine to thymine, WGBS libraries feature an extremely skewed nucleotide composition which can negatively affect Illumina sequencing. Consequently, WGBS reads showed substantially lower sequencing quality scores at cytosine/guanine base pairs compared to TAPS (Fig. 3B). To compensate for the nucleotide composition bias, at least 10 to 20% PhiX DNA (a base-balanced control library) is commonly added to WGBS libraries *(23)*. Accordingly, we supplemented the WGBS library with 15% PhiX. This, in combination with the reduced information content of BS-converted reads, and DNA degradation as a result of bisulfite treatment, resulted in significantly lower mapping rates for WGBS compared to TAPS (Fig. 3C and Table S5). Therefore, for the same sequencing cost (one NextSeq High Output run), the average depth of TAPS exceeded that of WGBS (24.4× and 13.1×, respectively; Table S6). Furthermore, TAPS resulted in fewer uncovered regions, and overall showed a more even coverage distribution, even after down-sampling to the same sequencing depth as WGBS (inter-quartile range: 7 and 11, respectively; Fig. 3D and Table S6). These results demonstrate that TAPS dramatically improved sequencing quality compared to WGBS, while effectively halving the sequencing cost.

The higher and more even genome coverage of TAPS resulted in a larger number of CpG sites covered by at least three reads. With TAPS, 91.8% of all 43,205,316 CpG sites in the mouse genome were covered at this level, compared to only 77.5% with WGBS (Fig. 4A and fig. S10). TAPS and WGBS resulted in highly correlated methylation measurements across chromosomal regions (Fig. 4D and fig. S11). TAPS slightly under-estimated the per-base modification rate, in line with the approx. 15% non-conversion rate for modified C (Fig. 3A). On a per-nucleotide basis, 32,610,160 CpG positions were covered by at least three reads in both methods (Fig. 4C). Within these sites, we defined “modified CpGs” as all CpG positions with a modification level of at least 10% *(24)*. Using this threshold, 95.5% of CpGs showed matching modification states between TAPS and WGBS. 98.2% of all CpGs that were covered by at least three reads and found modified in WGBS were recalled as modified by TAPS, indicating good agreement between WGBS and TAPS (Fig. 4B). When comparing modification levels, the fraction of modified reads per CpG, we observed good correlation between TAPS and WGBS (Pearson *r* = 0. 59, *p* < 2e-16, Fig. 4C). Notably, TAPS identified a subset of highly modified CpG positions which were missed by WGBS (Fig. 4C, bottom right corner). TAPS was thus able to identify DNA modifications in regions inaccessible by WGBS (Fig. 4E), spanning genes and CpG Islands (CGI). CGIs in particular were generally better covered by TAPS, even when controlling for differences in sequencing depth between WGBS and TAPS (Fig. 4F). Interestingly, TAPS identified higher average modification levels inside CGIs than WGBS, even though TAPS generally showed lower modification levels outside CGIs (fig. S12). This could suggest that CGI methylation levels are in fact higher than previously believed. Together, these results indicate that TAPS can directly replace WGBS, and in fact provides a more comprehensive view of the methylome than WGBS.

**Fig. 4.**
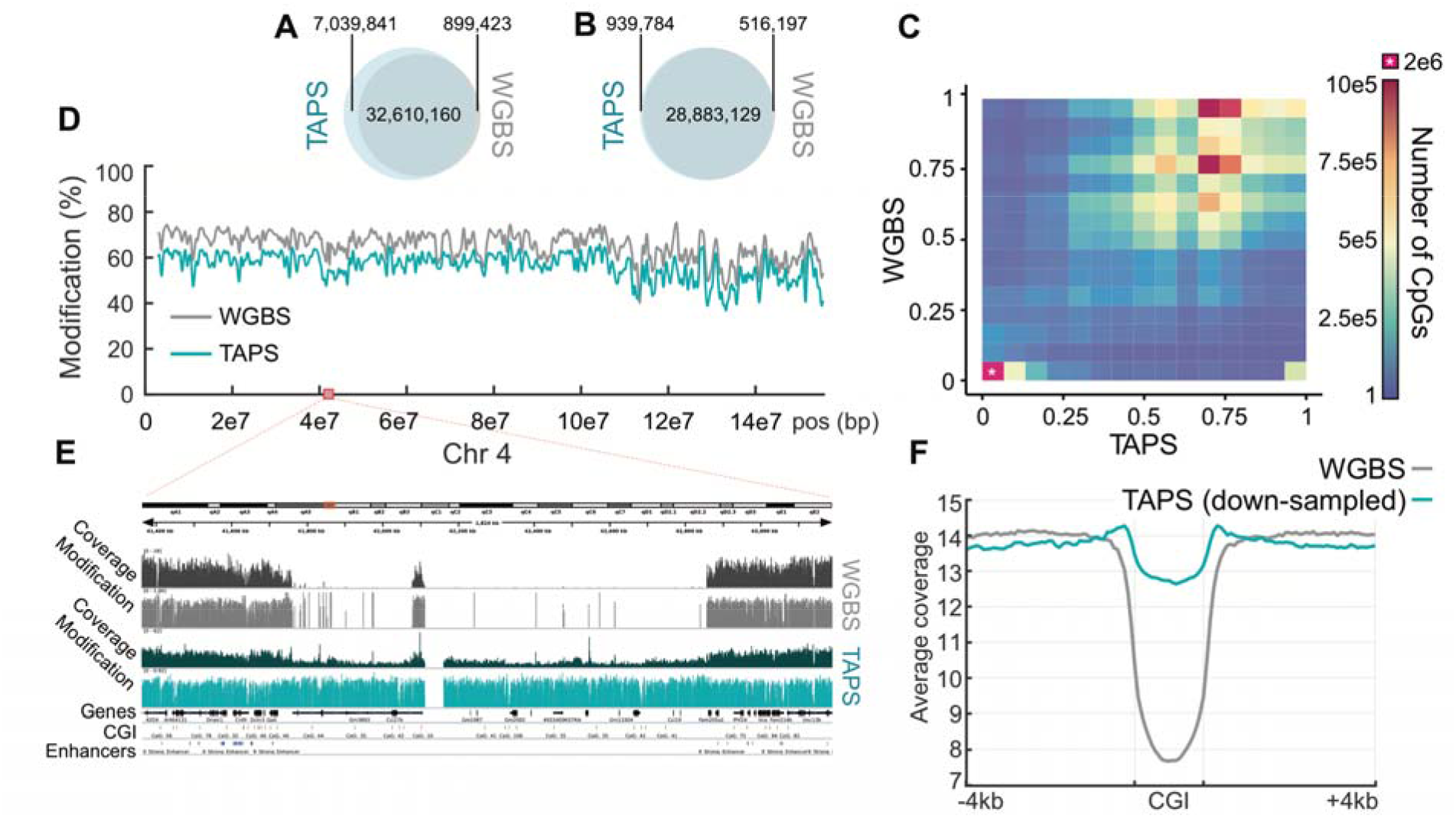
Comparison of genome-wide methylome measurements by TAPS and WGBS. **(A)** CpG sites covered by at least three reads by TAPS alone, both TAPS and WGBS, or WGBS alone. **(B)** Number of CpG sites covered by at least three reads and modification level > 0.1 detected by TAPS alone, TAPS and WGBS, or WGBS alone. **(C)** Heatmap representing the number of CpG sites covered by at least three reads in both TAPS and WGBS, broken down by modification levels as measured by each method. To improve contrast, the first bin, containing CpGs unmodified in both methods, was excluded from the color scale and is denoted by an asterisk. **(D)** Example of the chromosomal distribution of modification levels (in %) for TAPS and WGBS. Average fraction of modified CpGs per 100kb windows along mouse chromosome 4, smoothed using a Gaussian-weighted moving average filter with window size 10. **(E)** Example region on chromosome 4. TAPS provides information on regions that were not covered by WGBS, spanning both exons and CpG Islands (CGI). **(F)** Average sequencing coverage depth in all mouse CpG islands (binned into 20 windows) and 4kbp flanking regions (binned into 50 equally sized windows). To account for differences in sequencing depth, all mapped TAPS reads were down-sampled to match the mean coverage of WGBS across the genome.

Finally, we tested TAPS with low input DNA and showed that TAPS can work with down to 10 pg of gDNA, close to single-cell level. TAPS also works effectively with down to 1 ng of circulating cell-free DNA. These results demonstrate the potential of TAPS for low input DNA and clinical applications (fig. S13).

In summary, we have developed a series of PS-derived bisulfite-free, base-resolution sequencing methods for cytosine epigenetic modifications and demonstrated the utility of TAPS for whole-methylome sequencing. By using mild enzymatic and chemical reactions to detect 5mC and 5hmC directly at base-resolution without affecting unmodified cytosines, TAPS outperforms bisulfite sequencing in providing a high quality and more complete methylome at half the sequencing cost. As such TAPS could replace bisulfite sequencing as the new standard in DNA methylcytosine and hydroxymethylcytosine analysis. Rather than introducing a bulky modification on cytosine in the bisulfite-free 5fC sequencing method reported recently *(14, 15)*, TAPS converts modified cytosine into DHU, a near natural base, which can be “read” as T by common polymerases and is potentially compatible with PCR-free DNA sequencing. TAPS is compatible with a variety of downstream analyses, including but not limit to, pyrosequencing, methylation-sensitive PCR, restriction digestion, MALDI mass spectrometry, microarray and whole-genome sequencing. With further development, we expect TAPS to revolutionize DNA epigenetic analysis, and to have wide applications in academic research and clinical diagnostics, especially in sensitive low-input samples, such as circulating cell-free DNA (*25*) and single-cell *analysis (26, 27)*.

## Acknowledgements

We would like to acknowledge P. Spingardi, G. Berridge and B. Kessler for helping with the HPLC-MS/MS; P. Brennan and G.F. Ruda for helping with the NMR; F. Howe, S. Kriaucionis and C. Goding for critical reading of this manuscript

## Author contributions

Y.L. and C.-X.S. conceived the study and designed the experiments. Y.L. and P.S. performed the experiments with the help from F.Y., C.B., L.C. and Y.B. G.V. and B.S.-B. developed processing software. G.V., M.T. and B.S.-B. analyzed data. Y.L., B.S.-B. and C.-X.S. wrote the manuscript.

## Data and materials availability

Supplementary materials contain additional data. All data needed to evaluate the conclusions in the paper are present in the paper or the supplementary materials. All sequencing data are available through GEO no. GSE112520. The software used to process TAPS data can be downloaded from https://bitbucket.org/bsblabludwig/modificationfinder.

## Supplementary Materials

Materials and Methods

Figures S1-S13

Tables S1-S6

References (28-34)

